# Qinichelins, novel catecholate-hydroxamate siderophores synthesized via a multiplexed convergent biosynthesis pathway

**DOI:** 10.1101/163170

**Authors:** Jacob Gubbens, Changsheng Wu, Hua Zhu, Dmitri V. Filippov, Bogdan I. Florea, Sébastien Rigali, Herman S. Overkleeft, Gilles P. van Wezel

## Abstract

The explosive increase in genome sequencing and the advances in bioinformatic tools have revolutionized the rationale for natural product discovery from actinomycetes. In particular, this has revealed that actinomycete genomes contain numerous orphan gene clusters that have the potential to specify many yet unknown bioactive specialized metabolites, representing a huge unexploited pool of chemical diversity. Here, we describe the discovery of a novel group of catecholate-hydroxamate siderophores termed qinichelins (**2–5**) from *Streptomyces* sp. MBT76. Correlation between the metabolite levels and the protein expression profiles identified the biosynthetic gene cluster (BGC; named *qch*) most likely responsible for qinichelin biosynthesis. The structure of the molecules was elucidated by bioinformatics, mass spectrometry and NMR. Synthesis of the qinichelins requires the interplay between four gene clusters, for its synthesis and for precursor supply. This biosynthetic complexity provides new insights into the challenges scientists face when applying synthetic biology approaches for natural product discovery.

Pride repository reviewer account details:

URL: <u>https://www.ebi.ac.uk/pride/archive/login</u>

Project accession: PXD006577

Username

Password: 3H0iM1FK

## INTRODUCTION

Actinobacteria are renowned for their ability to manufacture a diversity of bioactive small molecules.^*1,2*^ The traditional approach for microbial natural product (NP) discovery typically involves high-throughput screening of crude extracts derived from cultivable actinomycetes, followed by iterative bioassay-guided fractionation and structure elucidation. This drug-discovery pipeline has rewarded us with many useful therapeutic agents, but also turned big pharma away from NPs for drug-discovery programs due to high cost and chemical redundancy.^*3,4*^ The explosive increase in genome-sequence information has uncovered a vast and yet untapped biosynthetic potential and metabolic diversity, which has brought the microbial NPs back into the spotlight. However, many of the biosynthetic gene clusters (BGCs) discovered by genome mining are poorly expressed under laboratory conditions, and a major new challenge lies in finding the triggers and cues to activate their expression.^*5*^ Such approaches include, among others, chemical triggers, microbial cocultivation, induction of antibiotic resistance, and heterologous gene expression. ^*6-10*^ In addition, the advances in genetic tools applied in synthetic biology, such as transformation-associated recombination (TAR), Red/ET recombination, and CRISPR-Cas9, had aided in the discovery of cryptic products through engineering of their biosynthetic pathways.^*11*^

A second bottleneck in genomics-based approaches is to establish a link between genomic and metabolomic data.^*5,12*^ It is difficult to assign the genetic basis for specific chemical scaffolds through bioinformatics analysis alone, largely due to nature’s flexibility in catalytic enzymology, i.e. enzyme promiscuity ^*13*^ and crosstalk among different gene clusters.^*14,15*^ The latter offers a significant hurdle in drug-discovery approaches that are based solely on heterologous expression of single gene clusters.^*16*^ This gap can be bridged by genomics-based methodologies that allow statistical correlation between transcript or protein expression levels on the one hand and abundance of the bioactive molecules on the other; such correlations allow the linkage between the biosynthetic genes and the bioactivity of interest, as we and others have previously exemplified. ^*17-20*^ Subsequent bioinformatics analysis of a biosynthetic gene cluster (BGC) provides important (partial) structural information, whereby the architectures of polyketides or nonribosomal peptides may be inferred from genetic organization due to assembly collinearity.^*21*^ This information can guide researchers to optimize compound isolation and identification, so as to recover sufficient quantity of targeted metabolites(s) from highly complex matrices to warrant *de novo* structural elucidation.^*22*^

Iron is an essential element required for a variety of metabolic processes in living organisms, but iron acquisition is challenging for most microorganisms due to the insolubility of iron(III). To cope with iron-limiting conditions, bacteria have evolved siderophores, which are synthesized by nonribosomal peptide synthases (NRPS) and act as iron scavengers.^*23*^ Siderophores commonly contain three bidentate ligands in one structure to coordinate iron by formation of a stable octahedral complex. Their chemical topologies and biosynthetic machineries have been studied extensively,^*14*, *15, 24*^ and a wide range of structures have been reported. Siderophores are generally classified into the following classes, according to the functionalities responsible for the chelation of ferric iron (Fe^3+^): catecholates, hydroxamates, (hydroxy)-carboxylates, and mixed-ligands thereof.^*23*^ Members of the mixed catecholate-hydroxamate sub-family,including rhodochelin,^*15*^ heterobactins,^*25*^rhodobactin,^*26*^ lystabactins,^*27*^ mirubactin,^*28*^ and S-213L,^*29*^ feature both a 2,3-dihydroxybenzoate(s) (2,3-DHB) moiety. And (modified) *δ-N*-hydroxyornithine residues within the same molecule. Consequently, the biosynthesis of catecholate-hydroxamate siderophores is always initiated by loading 2,3-DHB as starter unit into the modular NRPS assembly line, followed by successive incorporation of amino acids, including ornithine, into the growing peptide chain.^*15, 25, 28*^

*Streptomyces* sp. MBT76 was previously identified as a prolific producer of antibiotics against a number of ESKAPE pathogens under specific growth conditions,^*30*^ and time-course metabolomics analyses revealed it could produce a diversity of secondary metabolites, such as isocoumarins, prodiginines, acetyltryptamine, and fervenulin.^*31*^ More recently, the activation of a type II polyketide synthase (PKS) gene cluster (*qin*) in *Streptomyces* sp. MBT76 induced the production of many other polyketides, including a group of novel glycosylated pyranonaphthoquinones (qinimycins).^*32*^ As this metabolic spectrum was dominated by polyketides, while the BGCs for NRPS are as commonplace as PKS in bacterial genomes, ^*33, 34*^ we anticipated that peptides were underrepresented in our studies. Here we describe the discovery and characterization of qinichelins, new mixed-type catecholate-hydroxamate siderophores from *Streptomyces* sp. MBT76. The aforementioned limitations in genome-mining strategies were overcome through varying the growth conditions to fluctuate peptide production. Quantitative proteomics allowed the connection of the NRPS gene clusters to their metabolic products, enabling the elucidation of qinichelin and its BGC. This pipeline may also aid the discovery of other families of molecules.

## RESULTS AND DISCUSSION

### Biosynthetic loci for catechol-peptide siderophores are dispersed through the genome of *Streptomyces* sp. MBT76

Previously, AntiSMASH ^*35*^ analysis of the sequenced genome of *Streptomyces* sp. MBT76 identified 55 putative biosynthetic gene clusters (BGCs) specifying secondary metabolites.^*32*^ 16 of these contained gene(s) encoding NRPS, suggesting rich peptide metabolism. Our attention was in particular directed to three NRPS BGCs that lay well separated on the genome and governing the biosynthesis of catechol-peptide siderophores,. The first cluster was highly homologous to the *ent* cluster from *E. coli*,^*36*^ containing all the essential genes for enterobactin synthesis. Nevertheless, the gene organization in the *ent* cluster (*entA-C-E-B1-B2-F-D*) of *Streptomyces* sp. MBT76 differed from that in the *E. coli ent* cluster (*entA-F*). In *Streptomyces* sp. MBT76, the bifunctional *entB* gene was split into two separate genes *entB1* and *entB2,* encoding an isochorismatase and an acyl carrier protein (ACP) that are required for 2,3-DHB synthesis and loading, respectively (Figure 1b). This pattern was actually consistent with the *dhb* operon found in BGCs for the biosynthesis of the structurally related catechol-peptide siderophores griseobactin ^*37*^ and bacillibactin.^*38*^ Interestingly, a gene cluster (*gri*) for griseobactin synthesis lacking this exact *dhb* operon was also found in *Streptomyces* sp. MBT76 (Figure 1c), suggesting crosstalk between these two BGCs.

**Figure 1:**
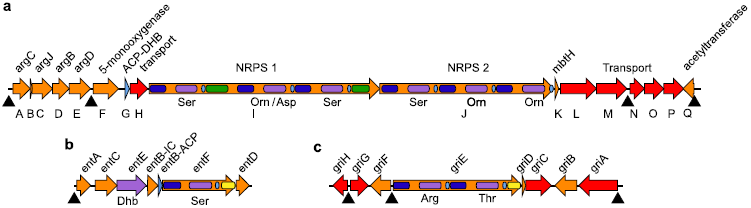
NRPS BGCs involved in catechol-type siderophore biosynthesis in *Streptomyces* sp. MBT76. BGCs for a potentially new siderophore (**a)**, for enterobactin **(b)**, and for griseobactin **(c)** could be identified. Carrier protein domains (ACP/PCP) are depicted in light blue, condensation domains in dark blue, epimerization domains in green, and thioesterase (termination) domains in yellow. Adenylation domains are shown in purple, together with their predicted substrates, and transport proteins in red. Triangles indicate the position of iron boxes likely bound by the iron repressor DmdR.

A third NRPS BGC that we designated *qch* (Table 1 and Figure 1a) contained *qchK* encoding an MbtH-like protein that is often found associated with NRPS BGCs,^*39*^ and multiple siderophore-related transporter genes *qchH* and *qchL-P*, suggesting the biosynthesis of a peptide siderophore. The presence of *qchG*, for an ACP homologous to the EntB-ACP domain, and the starter condensation (C) domain of *QchI*, which likely appends a DHB unit to the *N*-terminus of the peptide, indicated the presence of a DHB moiety in the final structure.^*36*^ However, genes required for DHB synthesis were not found within or near the *qch* cluster. Through phylogenetic analysis of adenylation (A) domains,^*40*^ the two core NRPS (QchI and QchJ) were predicted to produce a nonribosomal peptide with the sequence Ser-Orn(ornithine)/Asp-Ser-Ser-Orn-Orn, whereby no clear consensus prediction could be made for the second A domain. Two epimerization (E) domains in the first and third modules of QchI probably transform the stereochemistry of L-Ser into D-Ser, while the absence of an esterase (TE) domain at the terminus of QchJ indicated an unusual release of the mature peptide. Consistent with the A-domain analysis of the NRPS, *qch* contained four genes (*qchA* and *qchCDE*) highly similar to the *arg* genes *argC, argJ, argB,* and *argD* which are required for the synthesis of the nonproteinogenic amino acid ornithine from glutamate.^*41*^ Two accessory genes were predicted to be involved in tailoring of Orn: *qchF* coding for an L-ornithine-5-monooxygenase, and *qchQ* coding for a GCN5-related *N*-acetyltransferase.^*42*^ Interestingly, a canonical *arg* cluster ^*41*^ for arginine metabolism was also found in the *Streptomyces sp.* MBT76 genome, including the regulator *argR* and *argE-H* for the conversion of ornithine to arginine, all of which were lacking in the *qch* gene cluster.

**Table 1:**
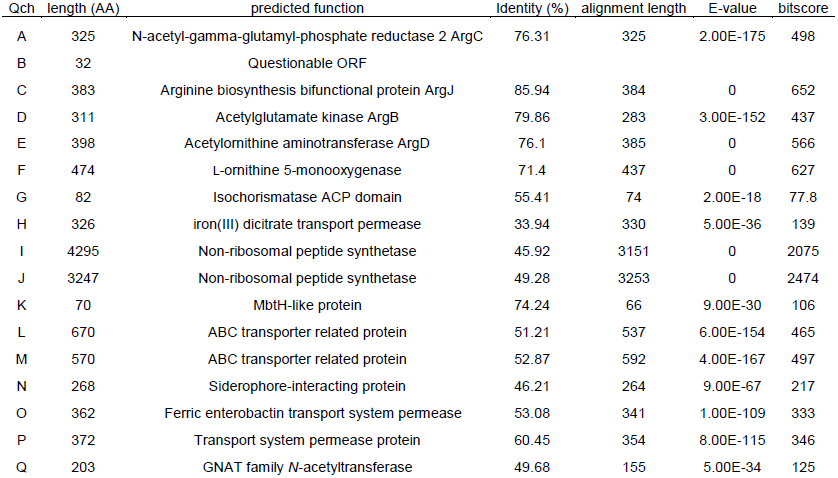
Blastp analysis of NRPS cluster

Taken together, bioinformatics analysis suggested: i) intertwined functional crosstalk among three separate BGCs in *Streptomyces* sp. MBT76, allowing production of catecholate-peptide siderophores; ii) that up to three types of catecholate-peptide siderophores might be produced by the strain, sharing one set of DHB genes; iii) potentially novel siderophores derived from the *qch* NRPS that encompass a connectivity of DHB-Ser-Orn/Asp-Ser-Ser-Orn-Orn, whereby the Orn-decorating genes could further diversify the final products. A search of the CAS database (American Chemical Society, <u>http://scifinder.cas.org</u>) using the predicted DHB-Ser-Orn-Ser-Ser-Orn-Orn sequence as a query, retrieved S213L (**1**, Figure 2),^*29*^ a patented antibiotic/antifungal siderophore with the sequence DHB-Ser-Orn-Ser-Orn-hOrn-chOrn, as closest hit. The partial sequence for the S213L BGC has been described,^*43*^ and the linear structure of S213L is compatible with the modular organization of our central NRPS QchI and QchJ, except for the specificity of the fourth A domain (Ser vs. Orn). Therefore, we speculated that the *qch* cluster might produce a related compound **2**, which contains a serine residue instead of ornithine as the fourth amino acid.

**Figure 2:**
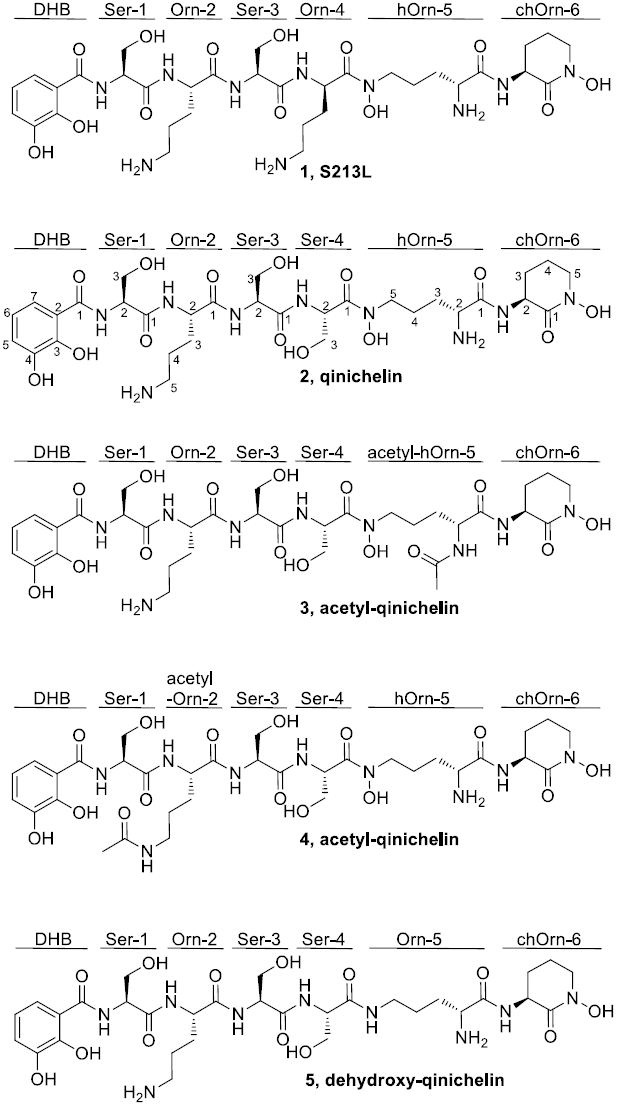
Molecular structures of S213L (1), qinichelin (2), acetyl-qinichelin (3), and dehydroxy-qinichelin (4). Abbreviations of moieties are shown to facilitate the comparison of respective structure. DHB, dihydroxybenzoate; Ser, serine; Orn, ornithine; hOrn, δ-*N*-hydroxy-ornithine; chOrn, cyclized δ-*N*-hydroxy-ornithine hydroxamate. Acetylation of qinichelin can occur on two positions as indicated. The shown configurations of DHB-D-Ser-L-Orn-D-Ser-L-Ser-L-hOrn-L-chOrn in qinichelins were deduced from biosynthetic considerations (see Figure 5).

### Proteomics analysis of the *qch* cluster and identification of the qinichelins

We previously described the *natural product proteomining* pipeline, which makes use of the strong correlation between the amount of a (bioactive) molecule produced and the expression level of its biosynthetic proteins.^*44*^ This was applied to efficiently connect genes (genotype) to a given metabolite or bioactivity of interest (chemotype). The reverse analysis whereby the expression level of a targeted BGC (known genotype) is used to predict its yet uncharacterized molecule that is produced (unknown chemotype), should be equally feasible. Accordingly, this reverse *proteomining* could complement a genome-mining strategy to facilitate the discovery of novel compounds.

As a prerequisite, sufficient fluctuation of protein levels should be achieved as a result of varying growth conditions.^*44*^ Accordingly, *Streptomyces* sp. MBT76 was grown in modified liquid minimal medium (NMMP), supplemented with (A) no additive (control), (B) 2% (w/v) NaCl, (C) 1% (w/v) starch, (D) 0.8% (w/v) peptone, or (E) 0.6% (w/v) yeast extract. Subsequent quantitative proteomics analysis of whole-cell lysates, using two mixtures of three samples to compare all growth conditions, yielded 1,472 protein identifications, wherein relative expression levels of 1,174 proteins were quantified with at least two independent events, including proteins belonging to the BGCs of interest (Table 2). Cultures grown in NMMP with peptone and, remarkably, in NMMP without additives, showed strong expression of the *qch* gene cluster, as demonstrated by the marked upregulation of QchF and QchH-J when compared to e.g. condition B (NMMP with 2% NaCl). Due to its small size, the dataset for the QchG protein contained only three quantifications, while the expression of QchA and ArgC could not be differentiated due to their high sequence similarity. However, the fluctuation pattern of QchA/ArgC for the five culture conditions was in line with that of all the other detected Arg proteins, strongly suggesting that the observed signals for QchA/ArgC were most likely dominated by ArgC rather than QchA.

**Table 2:** Quantitative proteomics analysis

| proteins | normalized expression ratio (<sup>2</sup>log)<sup>a</sup> |  |  |  |  |  | number of quantifications<sup>b</sup> |  |  |  |  |  |
| --- | --- | --- | --- | --- | --- | --- | --- | --- | --- | --- | --- | --- |
|  | D/A | B/A | B/D | D/C | E/C | E/D | D/A | B/A | B/D | D/C | E/C | E/D |
| **Qinichelin** |  |  |  |  |  |  |  |  |  |  |  |  |
| QchA/ArgC<sup>c</sup> | -1.6 | -0.8 | 1.0 | -1.7 | -2.2 | -0.3 | 7 | 7 | 7 | 3 | 3 | 3 |
| QchF | -0.6 | -2.4 | -2.3 | 0.1 | 0.4 | 0.0 | 8 | 8 | 8 | 7 | 7 | 7 |
| QchG |  |  |  | 1.7 | 1.2 | -0.7 | 1 | 1 | 1 | 3 | 3 | 3 |
| QchH | -1.2 | -2.5 | -1.2 | -1.0 | -1.5 | -1.2 | 6 | 6 | 6 | 4 | 4 | 4 |
| QchI | -0.2 | -3.1 | -3.3 | -0.7 | -1.1 | -0.5 | 12 | 12 | 12 | 7 | 7 | 7 |
| QchJ | -1.3 | -3.0 | -1.7 | -0.2 | -0.4 | -1.0 | 22 | 22 | 22 | 13 | 13 | 13 |
| **Enterobactin** |  |  |  |  |  |  |  |  |  |  |  |  |
| EntA | -2.1 | -0.6 | 1.3 |  |  |  | 3 | 3 | 3 | 0 | 0 | 0 |
| EntB-IC | -0.8 | 0.9 | 1.1 |  |  |  | 3 | 3 | 3 | 1 | 1 | 1 |
| **Griseobactin** |  |  |  |  |  |  |  |  |  |  |  |  |
| GriG | -1.3 | -1.5 | 0.1 | -2.2 | -2.2 | -0.7 | 8 | 8 | 8 | 8 | 7 | 7 |
| GriF |  |  |  | -1.4 | -1.2 | -0.1 | 0 | 0 | 0 | 6 | 6 | 6 |
| GriE | -0.8 | -1.4 | -0.3 |  |  |  | 3 | 3 | 3 | 0 | 0 | 0 |
| **Arginine biosynthesis cluster** |  |  |  |  |  |  |  |  |  |  |  |  |
| QchA /ArgC<sup>c</sup> | -1.6 | -0.8 | 1.0 | -1.7 | -2.2 | -0.3 | 7 | 7 | 7 | 3 | 3 | 3 |
| ArgJ | -0.9 | -0.9 | 0.0 | -0.6 | -0.8 | -0.9 | 7 | 7 | 7 | 6 | 5 | 5 |
| ArgD | -1.0 | -0.2 | 0.7 | -1.7 | -0.5 | 0.9 | 2 | 2 | 2 | 2 | 2 | 2 |
| ArgR | -1.5 | 0.1 | 1.5 | -2.5 | -1.0 | 1.1 | 9 | 9 | 9 | 4 | 4 | 4 |
| ArgG | -0.6 | -0.6 | -0.1 | -0.6 | -0.7 | 0.0 | 4 | 4 | 4 | 2 | 2 | 2 |
| ArgH | -1.5 | -1.1 | 0.6 | -1.7 | -0.8 | 0.5 | 8 | 8 | 8 | 9 | 9 | 9 |

<sup>a</sup> Changes in protein expression levels observed for the indicated proteins when compared among growth conditions A-E. <sup>b</sup> Number of quantifications events used to calculate the expression ratios. Quantifications based on less than two events (italicized) were discarded. <sup>c</sup> The sequences of QchA and ArgC were very similar, resulting in insufficient unique peptides for quantification. Instead, quantification is shown for non-unique peptides.

The proteomics analysis demonstrated the expression of the *qch* cluster in, amongst others, culture condition D (NMMP with peptone) and thus indicated the existence of the corresponding catecholate-peptide siderophore under these growth conditions. In our previous metabolomics study of *Streptomyces* sp. MBT76 under the same conditions,^*31*^ no siderophores were identified, which is most likely due to the use of ethyl acetate for the extraction, which is not suited for the isolation of the hydrophilic peptidic siderophores. Therefore, here spent media from five culturing conditions were desalted only and directly subjected to reverse-phase LC-MS analysis (in positive mode) without any prior extraction, resulting in the detection of a signal at *m*/*z* 772.3 for NMMP (A) and NMMP with peptone (D), with the strongest signal obtained for A (Figure 3a). The fluctuation pattern of this molecule correlated well with the expression level of the *qch* gene cluster, suggesting this may be the sought-after compound **2**. In addition to the molecular ion [M + H]^+^ at *m*/*z* 772.3 for the iron-free compound, a co-eluting peak was observed at *m*/*z* 825.3 corresponding to the iron-bound [M + Fe^3+^ - 2H]^+^ species. Figure 3a depicts the combined signals for both species to compensate for any differences in iron(III) concentration among the different culture conditions.

**Figure 3:**
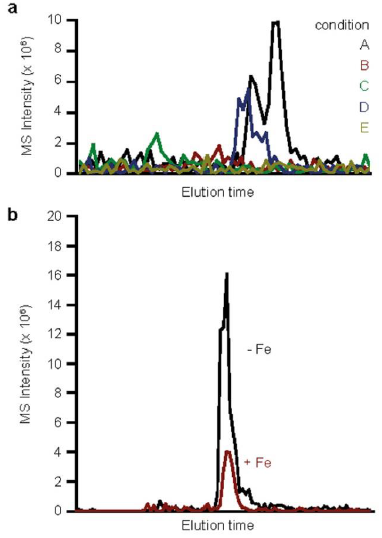
Comparison of qinichelin production by LC-MS analysis. Spent medium samples of *Streptomyces* sp. MBT76 grown in conditions A-E **(a)**; and in condition A in the absence (red line) or presence (black line) of Fe^3+^ (**b**), respectively, were compared. Shown are summed traces of [M+H]^+^ 772.3 *m*/*z* and [M+Fe^3+^-2H]^+^ 825.3 *m*/*z* ± 0.5 Da.

To confirm the structure of **2**, the spent medium of condition A was reanalyzed on a high resolution LTQ-orbitrap instrument, including both MS^1^ and MS^2^ analysis. Due to the use of formic acid instead of trifluoroacetic acid in the eluent, the MS^1^ spectrum of **2** presented the highest intensity at *m*/*z* 386.6794 assignable to [M + 2H]^2+^ species, followed by the [M + H]^+^ peak at *m*/*z* 772.3500 (Figure S1), within 0.5 ppm accuracy from the predicted mass. Indeed, the MS^2^ analysis yielded almost all the expected fragmentation products of the predicted compound **2**, with complete sequence coverage for both the b- and y-ion series (Figure 4a). Moreover, the MS^2^ analysis corroborated the hydroxylation of two ornithines (hOrn-5 and chOrn-6) at the *C*-terminus, and the cyclization of the last ornithine (chOrn-6). The most intensive signals were obtained for the b5 and y2 ions, indicating that a potential hydroxamate bond might be more susceptible to cleavage than an amide bond. However, it was noteworthy that MS/MS analysis alone was not enough to indicate the presence of a peptide or isopeptide bond between Ser-4 and hOrn-5. To clarify this, the *m*/*z* 772.3 was used as a probe to guide the separation of target compound from the spent medium of condition A on reversed phase HPLC. The obtained semi-purified compound **2** was analyzed by ^1^H NMR (850 MHz, in D_2_O, Table 3), COSY, HSQC, and HMBC techniques (Figure S2–S6), which indeed supported a catecholate-hexapeptide architecture comprising three serine and three ornithine residues. In particular, a key HMBC correlation from H_2_-5 of hOrn-5 to C-1 of Ser-4 established that the linkage between these two residues was through the δ-hydroxylated-amine rather than α-amine of hOrn-5. The free amine group at C-2 of hOrn-5 could be also reflected by the upfield shifted H-2 (*δ*H 3.99), in contrast to the amidated H-2 of Orn-2 (*δ*H 4.44) and chOrn-6 (*δ*_H_ 4.40).

**Table 3:**
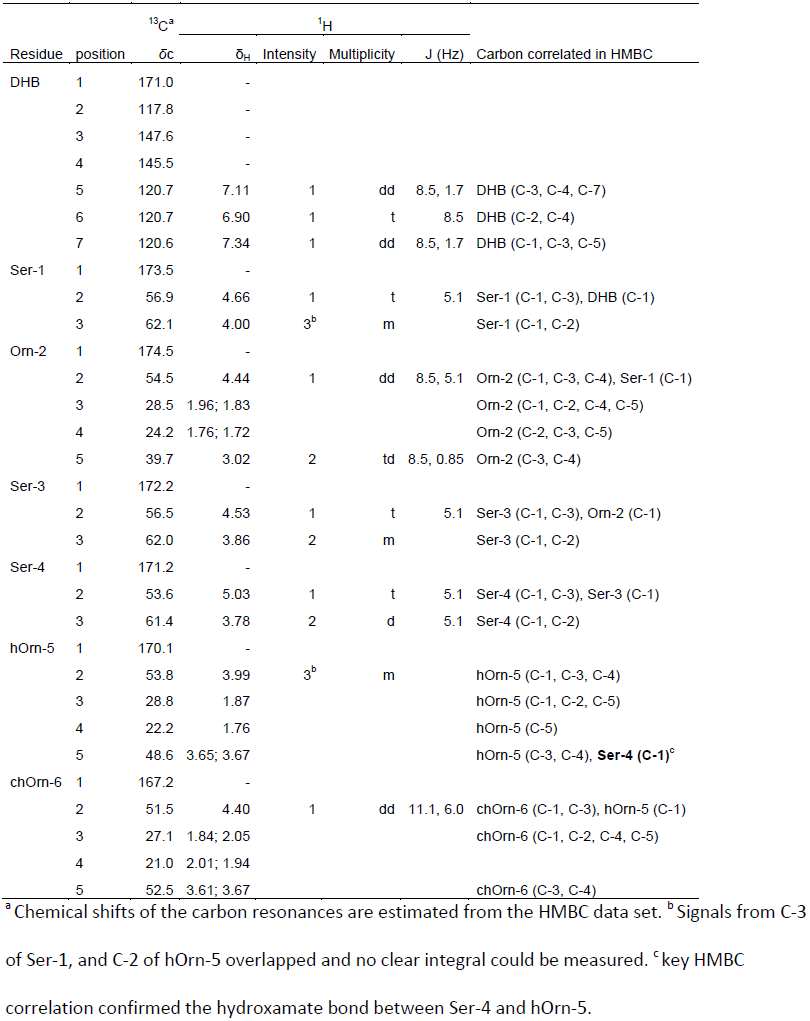
NMR data assignment of qinichelin (2) in D_2_O

**Figure 4:**
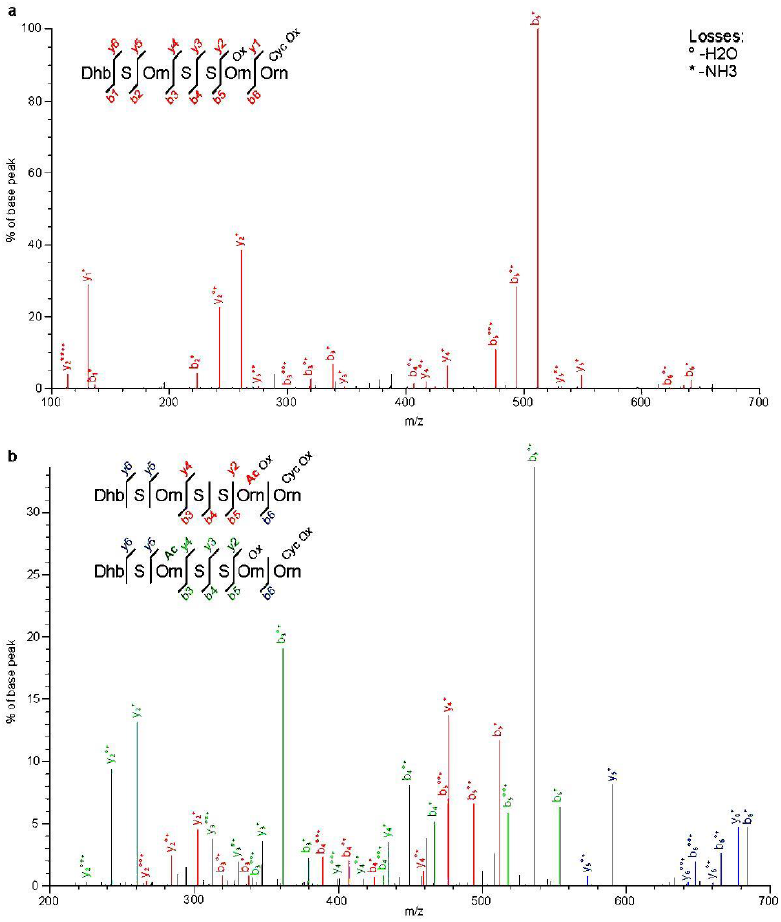
high resolution MS/MS analysis of qinichelin. A spent medium sample of *Streptomyces* sp. MBT76 grown in condition A was subjected to high resolution LC-MS/MS analysis to obtain insights into the structures of unacetylated (**a**) and acetylated (**b**) qinichelin.

Together, these experiments confirmed to existence and the precise chemical structure of compound **2**. With three iron-coordinating groups including one DHB moiety and two hydroxamates, our new compound resembles other mixed-ligand siderophores like amychelin ^*24*^ and gobichelin.^*45*^ This strongly suggested that compound **2** was a siderophore, which was named qinichelin. The name refers to the origin of *Streptomyces* sp. MBT76, which was isolated from the Qinling Mountains in China.^*30*^

### High resolution MS/MS analysis reveals production of qinichelin variants (3–5), griseobactin, but not enterobactin

We suspected that an acetylated analogue of qinichelin could be produced by *Streptomyces* sp. MBT76, because acetylation by an *N*-acetyltransferase encoded by *qchQ* had not yet been found in qinichelin. Indeed, we observed an [M + H]^+^ species at *m/z* 814.3587 for acetylated qinichelin, with a slightly longer retention time than qinichelin. The high abundance of an [M + H]^+^ species instead of [M + 2H]^2+^ already indicated that one of the two free amines in qinichelin, δ-NH_2_ in Orn-2 or α-NH_2_ in hOrn-5, was acetylated, while a derivative with both acetylations was not detected. Upon fragmentation for MS/MS analysis, a surprising result was obtained because the fragmentation spectrum (Figure 4b) corresponded to a mixture of two different acetylated peptides **3** and **4** (Figure 2). Some masses could only be assigned to acetylation at δ-NH_2_ in Orn-2 while other masses indicated acetylation of α-NH_2_ in hOrn-5. Since fragmentation of this [M + H]^+^ ion was less efficient than the unacetylated [M + 2H]^2+^ ion (Figure 4a), a complete sequence coverage could not be achieved for b- and y-ions. However, at least one b- or y-ion was present for each peptide/hydroxamate bond for both variants, thus providing strong evidence for the position of the posttranslational modification. In addition, qinichelin variant **5** gave a [M]^+^ peak at *m/z* 755.3314, and the characteristic fragment at *m/z* 512.2096 indicated an Orn-5 instead of an hOrn-5 residue (Figure S7). We did not obtain sufficient amounts of compounds **3–5** for 2D NMR analysis, as they are minor relative to **2**.

Since the proteomics analysis also revealed expression of the *ent* and *gri* clusters (Table 2), we attempted to find their respective products, enterobactin and griseobactin, by MS/MS analysis. Indeed, griseobactin could be readily detected with highest intensity at *m*/*z* 394.1720 for the [M+3H]^3+^ species, within 0.5 ppm of the expected mass. Another signal was observed for the [M+2H]^2+^ species at *m*/*z* 590.7538, with an MS/MS fragmentation pattern corresponding exactly with published data.^*46*^ Surprisingly, no enterobactin could be detected. This suggests that the *ent* cluster may only code for 2,3-DHB synthesis for griseobactin and qinichelin production in *Streptomyces* sp. MBT76, leaving *entF* non-functional.

### Qinichelin production belongs to the iron homeostasis regulon

To support the iron-chelating function of qinichelin and its possible role in iron homeostasis of *Streptomyces* sp. MBT76, we searched for the occurrence of iron boxes within the qinichelin BGC. Iron boxes are *cis*-acting elements with a 19 bp palindromic consensus sequence TTAGGTTAGGCTAACCTAA that are bound by DmdR1, the global iron regulator in *Streptomyces* species.^*47*^ When sufficient iron is available, the DmdR1-Fe^2+^ complex binds to iron boxes and represses the expression of siderophore biosynthetic and importer genes.^*47*^ The dramatic reduction in qinichelin production in an iron-rich condition suggested that the expression of *qch* cluster would also be under the negative control of DmdR1 (Figure 3b). Indeed, four highly conserved iron boxes were found within the BGC: (i) upstream of the predicted pentacistronic operon *qchA-E* involved in ornithine synthesis from glutamate, (ii) upstream of *qchF* coding for the L-ornithine 5-monooxygenase, (iii) upstream of the tricistronic operon (*qchN-P*) predicted to be involved in qinichelin transport, and (iv) upstream of *qchQ* that encodes the predicted qinichelin *N*-acetyltransferase (Table S1). The iron box identified 109 nt upstream of *qchF* displayed the perfect palindromic sequence TTAGGTTAGGCTAACCTAA, which made it highly likely that the central NRPS genes of the *qch* cluster were regulated by DmdR1. Furthermore, the iron box upstream of the predicted qinichelin transporter system (*qch*N-P in Figure 1) was more conserved than most of iron boxes identified upstream of other siderophore uptake system genes present in the *Streptomyces* sp. MBT76 genome (Table S1). In addition, three iron boxes were identified in the *gri* cluster and one in the *ent* cluster (Figure 1, and Table S1), suggesting that siderophore production in *Streptomyces* sp. MBT76 is indeed under control of DmdR1.

Interestingly, scanning for ARG boxes (consensus sequence CCATGCATGCCCATTGCATA) that are bound by the Arginine repressor ArgR ^*48*^ revealed no reliable *cis*-acting sequences upstream of the *qchA-E* operon. Instead, the *argCJBDR* gene cluster outside the qinichelin biosynthetic cluster displayed the putative ARG box at position -87 nt upstream of *argC*. This suggests differential regulation of the arginine biosynthetic genes from primary metabolism and those involved in secondary metabolism.

### Biosynthesis of qinichelins relies on coordination between multiple BGCs

The theoretical analysis and the experimental identification of griseobactin and qinichelins, allowed us to postulate an intertwined model for the production of catecholate-peptide siderophores in *Streptomyces* sp. MBT76 (Figure 5). The chorismate pathway within the *ent* gene cluster provides the building block 2,3-DHB to the three NRPS EntF, GriE, and QchI-QchJ, for enterobactin, griseobactin, and qinichelin respectively. The 2,3-DHB moiety is activated by 2,3-dihydroxybenzoate-AMP ligase EntE and subsequently transferred to stand-alone aryl carrier proteins QchG or EntB2. As the necessary gene coding for the aryl carrier protein is lacking in the griseobactin BGC, this requirement could be remedied by either QchG or EntB2 to deliver the activated 2,3-DHB starter unit for GriE. The further mechanisms for NRPS assembly of enterobactin and griseobactin have been elaborated elsewhere.^*37,49*^ The coordinated expression of multiple NRPS gene clusters for siderophore production in *Streptomyces* sp. MBT76 is striking but not unprecedented. Similar functional crosstalk between different NRPS BGCs was demonstrated for the assembly of the siderophores erythrochelin in *Saccharopolyspora erythraea* ^*14*^ and rhodochelin in *Rhodococcus jostii* RHA1.^*15*^ Such crosstalk could enable structural diversity for siderophores on the basis of a limited number of biosynthetic genes, and thus confer an evolutionary advantage for the producing bacteria in terms of iron acquisition. In particular, it would be advantageous for one bacterium to evolve specific siderophore(s) for their own benefit, to compete with the “siderophore pirates” that use siderophores biosynthesized by other species.^*50*^ For example, the structurally novel amychelin produced by *Amycolatopsis* sp. AA4, seems to frustrate “siderophore piracy” of *Streptomyces coelicolor* by inhibiting its development.^*24*^

**Figure 5.**
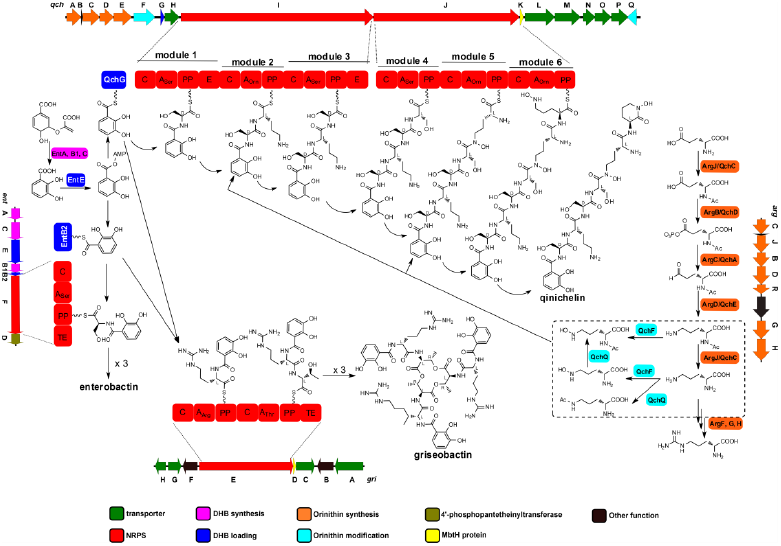
Intertwined biosynthetic pathways of catechol-peptide siderophores in *Streptomyces* sp. MBT76. The functional crosstalk among four entirely separate gene clusters ensures the assembly of two types of catechol-peptide siderophores, namely qinichelins and griseobactin. The DHB is supplied by *ent* gene cluster, which is shared by three NRPS systems. The DHB-hexapeptide backbone in qinichelin follows an orthodox colinear extension model, while the Orn building block could arise from either *qch* or canonical *arg* gene cluster. The differently modified Orn in the dashed boxes is likely accepted by the A_Orn_ domain of modules 2 and 5 to produce different qinichelin variants.

The assembly of the catecholate-hexapeptide backbone in qinichelin follows an orthodox linear logic of modular NRPS. Each module in QchI and QchJ contains an adenylation (A) domain for recognition of correct amino acid substrate, whereby Ser-1, Orn-2, Ser-3, Ser-4, hOrn-5, and hOrn-6, are sequentially bound and converted to aminoacyl adenylates. The two serine residues can be converted from the initial L-form ^*51*^ into its D-stereoisomer by the epimerization (E) domain in modules 1 and 3. After QchG-mediated incorporation of 2,3-DHB, each condensation (C) domain is successively used to elongate the chain by formation of a peptide bond with the activated amino acid, except for the isopeptide bond catalyzed by C domain 5, while the growing peptide chain is tethered to peptidyl carrier proteins (PP). Finally, qinichelin is released from the last PP domain through an intramolecular nucleophilic substitution of the δ-hydroxylamino group of L-hOrn-6 to the carbonyl group of the thioester. However, it is challenging to understand the enzymology responsible for this reaction, because a usual thioesterase (TE) domain (e.g. in NRPS assembling gobichelin ^*45*^ and heterobactin ^*25*^) required for peptide chain release is lacking in the C-terminus of QchJ. It is tempting to speculate that the C domain in module 6 catalyzes both the α-amidation of hOrn-6 to finalize the growing peptide chain, and δ-amidation to self-cyclize the last hydroxyornithine (chOrn-6) to release the peptide chain from NRPS system. A similar scenario for peptide chain release has recently been reported in the biosynthesis of scabichelin,^*52*^ a pentapeptide siderophore containing a C-terminal cyclic hydroxyornithine residue as in qinichelin.

The ornithine building block for qinichelin assembly may originate from either the *qch* cluster or from the canonical *arg* gene cluster,^*41*^ regulated by DmdR1 and ArgR, respectively. This would allow decoupling of qinichelin production from primary metabolism. The generated Orn precursor is further tailored, including hydroxylation at δ-NH_2_ by QchF, and/or acetylation at α-NH_2_ and δ-NH_2_ by QchQ. Alternatively, α-*N*-acetylation could arise from the bifunctional enzyme ArgJ (or its counterpart QchC) during ornithine precursor synthesis.^*53*^ The characterization of qinichelin congeners (**3**?**5**) provides evidence for substrate flexibility of the A_Orn_ domain in modules 2 and 5, whereby unmodified ornithine (Orn), δ-*N*-hydroxyl ornithine (hOrn), α-*N*-acetyl ornithine, δ-*N*-acetyl ornithine, and δ-*N*-hydroxyl-α-*N*-acetyl ornithine can be recognized and incorporated into the NRPS assembly. Still, we cannot rule out that QchQ post-translationally acetylates either free amine after construction of the final qinichelin. Indeed, it is difficult to discriminate between A domains activating Orn and/or hOrn through bioinformatics alone.^*54*^ However, since qinichelin (**2**) was the major chemical output of *qch* gene cluster, unmodified ornithine (Orn) and δ-*N*-hydroxyl ornithine (hOrn) are most likely preferred by A_Orn_ in module 2 and module 5, respectively.

## CONCLUSIONS

Actinomycetes adopt versatile strategies to biosynthesize structurally diverse secondary metabolites. This includes the production of a variety of siderophores, although it is not always clear what the advantage is in terms of the competition for iron in the environment. Functional crosstalk among multiple distantly located BGCs is not always predicted well by bioinformatics analysis. Therefore, chemical novelty may be missed if we solely rely on synthetic biology approaches, such as heterologous expression of a single BGC. The “protein-first” method, via reverse *natural product proteomining*, effectively identified the *qch* gene cluster expression in *Streptomyces* sp. MBT76, and further guided the characterization of qinichelins **2**–**5**, a family of new catecholate-hydroxamate siderophores. The principles presented in this work can be exploited to discover a broader range of chemical frameworks and to elucidate other intertwined biosynthetic scenarios.

### EXPERIMENTAL SECTION

### Strains and Growth conditions

*Streptomyces* sp. MBT76 isolation from Qinling mountain soil,^*30*^ general growth conditions, and genome sequencing (GenBank accession number: LNBE00000000) have been described before.^*31*^, ^*32*^ Here, *Streptomyces* sp. MBT76 was grown in liquid NMMP medium ^*55*^ containing 1% (w/v) glycerol and 0.5% (w/v) mannitol as carbon sources, but lacking polyethylene glycol. This basic NMMP medium were perturbed by using four different additives (or no additive) to create varying growth conditions: (A) no additive, (B) 2% (w/v)NaCl, (C) 1% (w/v) starch, (D) 0.8% (w/v) Bacto peptone (Difco), (E) 0.6% (w/v) Bacto yeast extract (Difco). For iron-starvation study, the minor element solution ^*55*^ was omitted from condition A. All the cultures of *Streptomyces* sp. MBT76 were incubated at 30 °C for 72 hours, with constant shaking at 220 rpm.

### Proteomics

*Streptomyces* sp. MBT76 cells were lysed using acetone/SDS as essentially described,^*56*^ since our ordinary ice sonication gave unsatisfactory results. Mycelium was washed in 2 ml of 100 mM Tris/HCl buffer (pH 7.5) supplemented with 10 mM MgCl_2_, and then resuspended in 5 ml of ice cold acetone for 10 min. After removal of acetone by centrifugation, the pellets were resuspended in 1 ml of 1% (w/v) SDS for 2 min. The resulting protein suspension (stored at -20 °C when necessary) was separated from the debris by centrifugation, and the protein concentrations were determined using the Qubit fluorometric system (Thermo) with BSA standard.

Some 167 μg of protein was precipitated for each sample using chloroform/methanol,^*57*^ and then dissolved using RapiGest SF surfactant (Waters, Milford MA). The proteins were further digested with trypsin after iodoacetamide treatment,^*58*^ and the resulting primary amines of the peptides were dimethyl labeled using three combinations of isotopomers of formaldehyde and cyanoborohydride on Sep-Pak C_18_ 200 mg columns (Waters, Milford MA), via CH_2_O + NaBH_3_CN, CD_2_O + NaBH_3_CN, and ^13^CD_2_O + NaBD_3_CN, as described.^*59*^ Light-, medium-, and heavy-labeled peptides with 4 Da mass differences were mixed 1:1:1 to obtain 0.5 mg for fractionation by cationic exchange (SCX) chromatography using a polysulfoethyl A column (PolyLC, 100 × 2.1 mm, particle size 5 mm, average pore size 200 Å). Mobile phases for SCX chromatography consisted of solvent A (10 mM KH_2_PO_4_, 20% (v/v) acetonitrile, Ph and solvent B (10 mM KH_2_PO_4_, 20% (v/v) acetonitrile, 0.5 M KCl, pH 3). The running program for SCX was a gradient of 0-18% solvent B in 18 CV (column volume), 18-30% solvent B in 6 CV, and 30-100% solvent B in 5 CV, at a constant flow rate of 250 μl/min. In total, 24 peptide fractions were collected for LC-MS/MS analysis on an LTQ-Orbitrap instrument (Thermo).^*58*^ Data analysis was performed using MaxQuant 1.4.1.2,^*60*^ whereby MS/ MS spectra were searched against a database of translated coding sequences obtained from genome of *Streptomyces* sp. MBT76. The mass spectrometry proteomics data have been deposited totheProteomeXchangeConsortium (<u>http://proteomecentral.proteomexchange.org</u>) via the PRIDE partner repository with the dataset identifier PXD006577.

### LC-MS analysis of metabolites

Spent medium samples were acidified with 1% (v/v) formic acid final concentration) and desalted using StageTips.^*61*^ 20 μl of samples were separated on an Finnigan Surveyor HPLC (Thermo) equipped with a Gemini C_18_ column (Phenomenex, 4.6 × 50 mm, particle size 3 mm, pore size 110 Å) at a flow rate of 1 mL/min and using a 0-50% B gradient in 10 CV. Mass spectrometry was performed using an Finnigan LCQ advantage (Thermo) equipped with an ESI source in the positive mode and scanning at 160 – 2,000 *m*/*z*.

For high resolution LC-MS/MS analysis on an LTQ-orbitrap the same setup was used as above for proteomics analysis,^*62*^ but using different run parameters. Mobile phases were: 0.1% (v/v) formic acid in H_2_O and B) 0.1% formic acid in acetonitrile. A 30 min 10-20% B gradient was followed by a 15 min 20-50% B gradient, both at a flow rate of 300 μL/min split to 250 nl/min by the LTQ divert valve. For each data-dependent cycle, one full MS scan (100-2,000 *m*/*z*) acquired at a resolution of 30,000 was followed by two MS/MS scans (100-2,000 *m*/*z*), again acquired in the orbitrap at a resolution of 30,000, with an ion selection threshold of 1 × 10^7^ counts but no charge exclusions. Other fragmentations parameters were as described for the proteomics analysis.^*62*^ After two fragmentations within 10 s, precursor ions were dynamically excluded for 120 s with an exclusion width of ± 10 ppm.

### Isolation of qinichelins

5 ml of spent medium from NMMP-grown cultures was desalted on Sep-Pak SPE C_18_ 200 mg columns (Waters). Columns were first washed with 1 ml of 80% (v/v) acetonitrile + 0.1% (v/v) formic acid, and then equilibrated with 1 ml of 0.1% (v/v) formic acid. 5 ml of spent medium was mixed with 1 ml of 5% (v/v) formic acid and loaded onto the column. After wash with 1 ml of 0.1% (v/v) formic acid, the column was eluted with 600 μl of 80% (v/v) acetonitrile + 0.1% (v/v) formic acid. The resulting sample was dried in a speedvac to remove acetonitrile and resuspended in 900 μl of 3% (v/v) acetonitrile + 0.1% (v/v) formic acid. This desalted sample was separated by HPLC on an Agilent 1200 series instrument equipped with a Gemini C_18_ column (Phenomenex, 250 × 10 mm, particle size 5 mm, pore size 110 Å), eluting with a gradient of acetonitrile in H_2_O adjusted with 0.15% (v/v) trifluoroacetic acid from 6% to 12%. The HPLC run was performed in 3 CV at a flow rate of 5 ml/min, and the fractions were collected based on UV absorption at 307 nm. All factions were analyzed by LC-MS (positive mode) to check the existence of the targeted mass at *m/z* 772.3. The fraction of interest was lyophilized, and subsequently reconstituted in deuterated water (D_2_O) for NMR (850 MHz) measurement.

### DmdR1 and ArgR regulon predictions

The putative binding sites for the iron utilization regulator DmdR1 and for the arginine biosynthesis regulator ArgR were detected on the chromosome of *Streptomyces* sp. MBT76 using the PREDetector software ^*63*^ and according to the method described.^*64*^ For the generation of the DmdR1 position weight matrix (PWM) we used the sequence of the iron box which lies at position -82 nt upstream of *desA* (SCO2782) and previously shown to be bound by DmdR1 in *S. coelicolor*.^*65*^ In order to acquire more highly reliable iron boxes to generate the PWM we scanned the upstream region of the orthologues of *desA* in five other *Streptomyces* species and retrieved their respective iron boxes (see supplementary Fig. S8). A set of ARG boxes experimentally validated in *S. clavuligerus* ^*66*^ and *S. coelicolor* ^*48*^ were used to generate the ArgR PWM (see supplementary Fig. S9).

## Acknowledgements

This work was supported by a grant from the Chinese Scholarship Council to CW and by a VENI grant from the Netherlands Foundation for Scientific Research (NWO) to GPvW.

## REFERENCES

1. Barka, E. A., Vatsa, P., Sanchez, L., Gavaut-Vaillant, N., Jacquard, C., Meier-Kolthoff, J., Klenk, H. P., Clément, C., Oudouch, Y., and van Wezel, G. P. (2016) Taxonomy, physiology, and natural products of the Actinobacteria, Microbiol Mol Biol Rev 80, 1– 43.

2. Bérdy, J. (2005) Bioactive microbial metabolites, J Antibiot (Tokyo) 58, 1–26.

3. Cooper, M. A., and Shlaes, D. (2011) Fix the antibiotics pipeline, Nature 472, 32.

4. Payne, D. J., Gwynn, M. N., Holmes, D. J., and Pompliano, D. L. (2007) Drugs for bad bugs: confronting the challenges of antibacterial discovery, Nat Rev Drug Discov 6, 29–40.

5. Doroghazi, J. R., Albright, J. C., Goering, A. W., Ju, K. S., Haines, R. R., Tchalukov, K. A., Labeda, D. P., Kelleher, N. L., and Metcalf, W. W. (2014) A roadmap for natural product discovery based on large-scale genomics and metabolomics,Nat Chem Biol 10, 963–968.

6. Craney, A., Ozimok, C., Pimentel-Elardo, S. M., Capretta, A., and Nodwell, J. R. (2012) Chemical perturbation of secondary metabolism demonstrates important links to primary metabolism, Chem Biol 19, 1020–1027.

7. Hosaka, T., Ohnishi-Kameyama, M., Muramatsu, H., Murakami, K., Tsurumi, Y., Kodani, S., Yoshida, M., Fujie, A., and Ochi, K. (2009) Antibacterial discovery in actinomycetes strains with mutations in RNA polymerase or ribosomal protein S12, Nat Biotechnol 27, 462–464.

8. Rutledge, P. J., and Challis, G. L. (2015) Discovery of microbial natural products by activation of silent biosynthetic gene clusters, Nat Rev Microbiol 13, 509–523.

9. van der Meij, A., Worsley, S. F., Hutchings, M. I., and van wezel, G. P. (2017) Chemical ecology of antibiotic production by actinomycetes, FEMS Microbiol Rev, in press.

10. Zhu, H., Sandiford, S. K., and van Wezel, G. P. (2014) Triggers and cues that activate antibiotic production by actinomycetes, J Ind Microbiol Biotechnol 41, 371–386.

11. Seyedsayamdost, M. R., and Clardy, J. (2014) Natural Products and Synthetic Biology, ACS Synth. Biol. 3, 745–747.

12. Wu, C., Kim, H. K., van Wezel, G. P., and Choi, Y. H. (2015) Metabolomics in the natural products field - a gateway to novel antibiotics, Drug Discov Today Technol 13, 11–17.

13. Wu, C., Medema, M. H., Läkamp, R. M., Zhang, L., Dorrestein, P. C., Choi, Y. H., and van Wezel, G. P. (2016) Leucanicidin and Endophenasides Result from Methyl-Rhamnosylation by the Same Tailoring Enzymes in *Kitasatospora* sp. MBT66, ACS Chem. Biol. 11, 478–490.

14. Lazos, O., Tosin, M., Slusarczyk, A. L., Boakes, S., Cortés, J., Sidebottom, P. J., and Leadlay, P. F. (2010) Biosynthesis of the Putative Siderophore Erythrochelin Requires Unprecedented Crosstalk between Separate Nonribosomal Peptide Gene Clusters, Chem. Biol. 17, 160–173.

15. Bosello, M., Robbel, L., Linne, U., Xie, X., and Marahiel, M. A. (2011) Biosynthesis of the siderophore rhodochelin requires the coordinated expression of three independent gene clusters in *Rhodococcus jostii* RHA1, J. Am. Chem. Soc. 133, 4587–4595.

16. Kolter, R., and van Wezel, G. P. (2016) Goodbye to brute force in antibiotic discovery?, Nat Microbiol 1, 15020.

17. Bumpus, S. B., Evans, B. S., Thomas, P. M., Ntai, I., and Kelleher, N. L. (2009) A proteomics approach to discovering natural products and their biosynthetic pathways, Nat Biotechnol 27, 951–956.

18. Gubbens, J., Zhu, H., Girard, G., Song, L., Florea, B. I., Aston, P., Ichinose, K., Filippov, D. V., Choi, Y. H., Overkleeft, H. S., Challis, G. L., and van Wezel, G. P. (2014) Natural product proteomining, a quantitative proteomics platform, allows rapid discovery of biosynthetic gene clusters for different classes of natural products, Chem Biol 21, 707–718.

19. Kersten, R. D., Ziemert, N., Gonzalez, D. J., Duggan, B. M., Nizet, V., Dorrestein, P. C., and Moore, B. S. (2013) Glycogenomics as a mass spectrometry-guided genome-mining method for microbial glycosylated molecules, Proc Natl Acad Sci U S A 110, E4407–4416.

20. Meier, J. L., Niessen, S., Hoover, H. S., Foley, T. L., Cravatt, B. F., and Burkart, M. D. (2009) An orthogonal active site identification system (OASIS) for proteomic profiling of natural product biosynthesis, ACS Chem Biol 4, 948–957.

21. Tietz, J. I., and Mitchell, D. (2016) Using Genomics for Natural Product Structure Elucidation, Curr. Top. Med. Chem. 16, 1645–1694.

22. Jensen, P. R., Chavarria, K. L., Fenical, W., Moore, B. S., and Ziemert, N. (2014) Challenges and triumphs to genomics-based natural product discovery, J. Ind. Microbiol. Biotechnol. 41, 203–209.

23. Miethke, M., and Marahiel, M. A. (2007) Siderophore-Based Iron Acquisition and Pathogen Control, Microbiol. Mol. Biol. Rev. 71, 413–451.

24. Seyedsayamdost, M. R., Traxler, M. F., Zheng, S. L., Kolter, R., and Clardy, J. (2011) Structure and biosynthesis of amychelin, an unusual mixed-ligand siderophore from *amycolatopsis sp.* AA4, J. Am. Chem. Soc. 133, 11434–11437.

25. Bosello, M., Zeyadi, M., Kraas, F. I., Linne, U., Xie, X., and Marahiel, M. A. (2013) Structural characterization of the heterobactin siderophores from *rhodococcus erythropolis* PR4 and elucidation of their biosynthetic machinery, J. Nat. Prod. 76, 2282–2290.

26. Dhungana, S., Michalczyk, R., Boukhalfa, H., Lack, J. G., Koppisch, A. T., Fairlee, J. M., Johnson, M. T., Ruggiero, C. E., John, S. G., Cox, M. M., Browder, C. C., Forsythe, J. H., Vanderberg, L. A., Neu, M. P., and Hersman, L. E. (2007) Purification and characterization of rhodobactin: A mixed ligand siderophore from *Rhodococcus rhodochrous* strain OFS, BioMetals 20, 853–867.

27. Zane, H. K., and Butler, A. (2013) Isolation, structure elucidation, and iron-binding properties of lystabactins, siderophores isolated from a marine *Pseudoalteromonas* sp., J. Nat. Prod. 76, 648–654.

28. Giessen, T. W., Franke, K. B., Knappe, T. A., Kraas, F. I., Bosello, M., Xie, X., Linne, U., and Marahiel, M. A. (2012) Isolation, structure elucidation, and biosynthesis of an unusual hydroxamic acid ester-containing siderophore from actinosynnema mirum,J. Nat. Prod. 75, 905–914.

29. Owaku, K., Umashima, T., Matsugami, M., Goto, M., Nakajima, T., Ito, T., Ikuko, K., Nozawa, A., and Miki, T. (2000) Antifungal antibiotic S-213L manufacture with *Streptomyces*, Japan.

30. Zhu, H., Swierstra, J., Wu, C., Girard, G., Choi, Y. H., van Wamel, W., Sandiford, S. K., and van Wezel, G. P. (2014) Eliciting antibiotics active against the ESKAPE pathogens in a collection of actinomycetes isolated from mountain soils, Microbiology 160, 1714–1725.

31. Wu, C., Zhu, H., van Wezel, G. P., and Choi, Y. H. (2016) Metabolomics-guided analysis of isocoumarin production by *Streptomyces* species MBT76 and biotransformation of flavonoids and phenylpropanoids, Metabolomics 12, 90.

32. Wu, C., Du, C., Ichinose, K., Choi, Y. H., and van Wezel, G. P. (2016) The cryptic qin gene cluster of *Streptomyces* sp. MBT76 specifies *C*-glycosylpyranonaphthoquinones, J. Nat. Prod., In press.

33. Donadio, S., Monciardini, P., and Sosio, M. (2007) Polyketide synthases and nonribosomal peptide synthetases: the emerging view from bacterial genomics, Nat Prod Rep 24, 1073–1109.

34. Medema, M. H., and Kottmann, R., and Yilmaz, P., and Cummings, M., and Biggins, J. B., and Blin, K., and de Bruijn, I., and Chooi, Y. H., and Claesen, J., and Coates, R.C., and Cruz-Morales, P., and Duddela, S., and Dusterhus, S., and Edwards, D. J., and Fewer, D. P., and Garg, N., and Geiger, C., and Gomez-Escribano, J. P., and Greule, A., and Hadjithomas, M., and Haines, A. S., and Helfrich, E. J., and Hillwig,M. L., and Ishida, K., and Jones, A. C., and Jones, C. S., and Jungmann, K., and Kegler, C., and Kim, H. U., and Kotter, P., and Krug, D., and Masschelein, J., and Melnik, A. V., and Mantovani, S. M., and Monroe, E. A., and Moore, M., and Moss, N., and Nutzmann, H. W., and Pan, G., and Pati, A., and Petras, D., and Reen, F. J., and Rosconi, F., and Rui, Z., and Tian, Z., and Tobias, N. J., and Tsunematsu, Y., and Wiemann, P., and Wyckoff, E., and Yan, X., and Yim, G., and Yu, F., and Xie, Y., and Aigle, B., and Apel, A. K., and Balibar, C. J., and Balskus, E. P., and Barona-Gomez, F., and Bechthold, A., and Bode, H. B., and Borriss, R., and Brady, S. F., and Brakhage, A. A., and Caffrey, P., and Cheng, Y. Q., and Clardy, J., and Cox, R. J., and De Mot, R., and Donadio, S., and Donia, M. S., and van der Donk, W. A., and Dorrestein, P. C., and Doyle, S., and Driessen, A. J., and Ehling-Schulz, M., and Entian, K. D., and Fischbach, M. A., and Gerwick, L., and Gerwick, W. H., and Gross, H., and Gust, B., and Hertweck, C., and Hofte, M., and Jensen, S. E., and Ju, J., and Katz, L., and Kaysser, L., and Klassen, J. L., and Keller, N. P., and Kormanec, J., and Kuipers, O. P., and Kuzuyama, T., and Kyrpides, N. C., and Kwon, H. J., and Lautru, S., and Lavigne, R., and Lee, C. Y., and Linquan, B., and Liu, X., and Liu, W., and Luzhetskyy, A., and Mahmud, T., and Mast, Y., and Mendez, C., and Metsa-Ketela, M., and Micklefield, J., and Mitchell, D. A., and Moore, B. S., and Moreira, L. M., and Muller, R., and Neilan, B. A., and Nett, M., and Nielsen, J., and O’Gara, F., and Oikawa, H., and Osbourn, A., and Osburne, M. S., and Ostash, B., and Payne, S. M., and Pernodet, J. L., and Petricek, M., and Piel, J., and Ploux, O., and Raaijmakers, J. M., and Salas, J. A., and Schmitt, E. K., and Scott, B., and Seipke, R. F., and Shen, B., and Sherman, D. H., and Sivonen, K., and Smanski, M. J., and Sosio, M., and Stegmann, E., and Sussmuth, R. D., and Tahlan, K., and Thomas, C. M., and Tang, Y., and Truman, A. W., and Viaud, M., and Walton, J. D., and Walsh, C. T., and Weber, T., and van Wezel, G. P., and Wilkinson, B., and Willey, J. M., and Wohlleben, W., and Wright, G. D., and Ziemert, N., and Zhang, C., and Zotchev, S. B., and Breitling, R., and Takano, E., and Glockner, F. O. (2015) Minimum Information about a Biosynthetic Gene cluster, Nat Chem Biol 11, 625–631.

35. Blin, K., Medema, M.H., Kazempour, D., Fischbach, M. A., Breitling, R., Takano, E., and Weber, T. (2013) antiSMASH 2.0-a versatile platform for genome mining of secondary metabolite producers., Nucleic Acids Res. 41, 1–9.

36. Gehring, A. M., Bradley, K. A., and Walsh, C. T. (1997) Enterobactin biosynthesis in Escherichia coli: isochorismate lyase (EntB) is a bifunctional enzyme that is phosphopantetheinylated by EntD and then acylated by EntE using ATP and 2,3- dihydroxybenzoate, Biochemistry (Mosc.) 36, 8495–8503.

37. Patzer, S. I., and Braun, V. (2010) Gene cluster involved in the biosynthesis of griseobactin, a catechol-peptide siderophore of *Streptomyces* sp. ATCC 700974, J. Bacteriol. 192, 426–435.

38. May, J. J., Wendrich, T. M., and Marahiel, M. A. (2001) The *dhb* Operon of *Bacillus subtilis* Encodes the Biosynthetic Template for the Catecholic Siderophore 2,3-Dihydroxybenzoate-Glycine-Threonine Trimeric Ester Bacillibactin, J. Biol. Chem. 276, 7209–7217.

39. Baltz, R. H. (2014) MbtH homology codes to identify gifted microbes for genome mining, J. Ind. Microbiol. Biotechnol. 41, 357–369.

40. Challis, G. L., Ravel, J., and Townsend, C. A. (2000) Predictive, structure-based model of amino acid recognition by nonribosomal peptide synthetase adenylation domains, Chem. Biol. 7, 211–224.

41. Rodríguez-García, A., de la Fuente, Á., Pérez-Redondo, R., Martín, J. F., and Liras, P. (2000) Characterization and expression of the arginine biosynthesis gene cluster of Streptomyces clavuligerus.,J. Mol. Microbiol. Biotechnol. 2, 543–550.

42. Vetting, M. W., Luiz, L. P., Yu, M., Hegde, S. S., Magnet, S., Roderick, S. L., and Blanchard, J. S. (2005) Structure and functions of the GNAT superfamily of acetyltransferases, Arch. Biochem. Biophys. 433, 212–226.

43. Chen, Y., Ntai, I., Ju, K. S., Unger, M., Zamdborg, L., Robinson, S. J., Doroghazi, J. R., Labeda, D. P., Metcalf, W. W., and Kelleher, N. L. (2012) A proteomic survey of nonribosomal peptide and polyketide biosynthesis in actinobacteria, J. Proteome Res. 11, 85–94.

44. Gubbens, J., Zhu, H., Girard, G., Song, L., Florea, B. I., Aston, P., Ichinose, K., Filippov, D. V., Choi, Y. H., Overkleeft, H. S., Challis, G. L., and van Wezel, G. P. (2014) Natural Product Proteomining, a Quantitative Proteomics Platform, Allows Rapid Discovery of Biosynthetic Gene Clusters for Different Classes of Natural Products., Chem. Biol. 21, 707–718.

45. Chen, Y., Unger, M., Ntai, I., McClure, R. A., Albright, J. C., Thomson, R. J., and Kelleher, N. L. (2013) Gobichelin A and B: Mixed-Ligand Siderophores Discovered Using Proteomics., MedChemComm 4, 233–238.

46. Albright, J. C., Goering, A. W., Doroghazi, J. R., Metcalf, W. W., and Kelleher, N. L. (2014) Strain-specific proteogenomics accelerates the discovery of natural products via their biosynthetic pathways, J Ind Microbiol Biotechnol 41, 451–459.

47. Flores, F. J., and Martín, J. F. (2004) Iron-regulatory proteins DmdR1 and DmdR2 of *Streptomyces coelicolor* form two different DNA-protein complexes with iron boxes, Biochem. J. 380, 497–503.

48. Pérez-Redondo, R., Rodríguez-García, A., Botas, A., Santamarta, I., Martín, J. F., and Liras, P. (2012) ArgR of *Streptomyces coelicolor* is a versatile regulator, PloS one 7, e32697.

49. Gehring, A. M., Bradley, K. A., and Walsh, C. T. (1997) Enterobactin biosynthesis in *Escherichia coli:* Isochorismate lyase (EntB) is a bifunctional enzyme that is phosphopantetheinylated by EntD and then acylated by ente using ATP and 2,3-dihydroxybenzoate, Biochemistry 36, 8495–8503.

50. Schubert, S., Fischer, D., and Heesemann, J. (1999) Ferric enterochelin transport in Yersinia enterocolitica: molecular and evolutionary aspects., J. Bacteriol. 181, 6387– 6395.

51. Lautru, S., and Challis, G. L. (2004) Substrate recognition by nonribosomal peptide synthetase multi-enzymes, Microbiology 150, 1629–1636.

52. Kodani, S., Bicz, J., Song, L., Deeth, R. J., Ohnishi-Kameyama, M., Yoshida, M., Ochi, K., and Challis, G. L. (2013) Structure and biosynthesis of scabichelin, a novel tris-hydroxamate siderophore produced by the plant pathogen *Streptomyces scabies* 87.22, Org. Biomol. Chem. 11, 4686–4694.

53. Rodriguez-Garcia, A., de la Fuente, A., Perez-Redondo, R., Martin, J. F., and Liras, P. (2000) Characterization and expression of the arginine biosynthesis gene cluster of Streptomyces clavuligerus, J. Mol. Microbiol. Biotechnol. 2, 543–550.

54. Stachelhaus, T., Mootz, H. D., and Marahiel, M. A. (1999) The specificity-conferring code of adenylation domains in nonribosomal peptide synthetases, Chem. Biol. 6, 493–505.

55. Kieser, T., Bibb, M. J., Buttner, M. J., Chater, K. F. & Hopwood, D. A. (2000) Practical Streptomyces genetics, John Innes Foundation, Norwich, UK.

56. Bhaduri, S., and Demchick, P. H. (1983) Simple and rapid method for disruption of bacteria for protein studies, Appl. Environ. Microbiol. 46, 941–943.

57. Wessel, D., and Flügge, U. I. (1984) A method for the quantitative recovery of protein in dilute solution in the presence of detergents and lipids, Anal. Biochem. 138, 141–143.

58. Gubbens, J., Janus, M., Florea, B. I., Overkleeft, H. S., and van Wezel, G. P. (2012) Identification of glucose kinase-dependent and -independent pathways for carbon control of primary metabolism, development and antibiotic production in *Streptomyces coelicolor* by quantitative proteomics., Mol. Microbiol. 86, 1490–1507.

59. Boersema, P. J., Raijmakers, R., Lemeer, S., Mohammed, S., and Heck, A. J. R. (2009) Multiplex peptide stable isotope dimethyl labelling for quantitative proteomics, Nat. Protoc. 4, 484–494.

60. Cox, J., and Mann, M. (2008) MaxQuant enables high peptide identification rates, individualized p.p.b.-range mass accuracies and proteome-wide protein quantification., Nat. Biotechnol. 26, 1367–1372.

61. Rappsilber, J., Mann, M., and Ishihama, Y. (2007) Protocol for micro-purification, enrichment, pre-fractionation and storage of peptides for proteomics using StageTips, Nat Protoc 2, 1896–1906.

62. Gubbens, J., Janus, M., Florea, B. I., Overkleeft, H. S., and van Wezel, G. P. (2012) Identification of glucose kinase-dependent and -independent pathways for carbon control of primary metabolism, development and antibiotic production in Streptomyces coelicolor by quantitative proteomics, Mol. Microbiol. 86, 1490–1507.

63. Hiard, S., Marée, R., Colson, S., Hoskisson, P. A., Titgemeyer, F., van Wezel, G. P., Joris, B., Wehenkel, L., and Rigali, S. (2007) PREDetector: A new tool to identify regulatory elements in bacterial genomes, Biochem. Biophys. Res. Commun. 357, 861–864.

64. Rigali, S., Nivelle, R., and Tocquin, P. (2015) On the necessity and biological significance of threshold-free regulon prediction outputs, Mol. BioSyst. 11, 333–337.

65. Tunca, S., Barreiro, C., Sola-Landa, A., Coque, J. J. R., and Martín, J. F. (2007) Transcriptional regulation of the desferrioxamine gene cluster of *Streptomyces coelicolor* is mediated by binding of DmdR1 to an iron box in the promoter of the desA gene, FEBS J. 274, 1110–1122.

66. Rodriguez-Garcia, A., Ludovice, M., Martin, J. F., and Liras, P. (1997) Arginine boxes and the argR gene in *Streptomyces clavuligerus*: evidence for a clear regulation of the arginine pathway, Mol Microbiol 25, 219–228.

